# A discrete approach to the external branches of a Kingman coalescent tree. Theoretical results and practical applications

**DOI:** 10.1101/818088

**Authors:** Filippo Disanto, Thomas Wiehe

**Affiliations:** Dipartimento di Matematica, Università di Pisa, Italy; Institut für Genetik, Universität zu Köln, Germany.

**Keywords:** Yule histories, Coalescent trees, External branches, Branch length, Combinatorics

## Abstract

The Kingman coalescent process is a classical model of gene genealogies in population genetics. It generates Yule-distributed, binary ranked tree topologies—also called *histories*—with a finite number of *n* leaves, together with *n* −1 exponentially distributed time lengths: one for each each layer of the history. Using a discrete approach, we study the lengths of the external branches of Yule distributed histories, where the length of an external branch is defined as the rank of its parent node. We study the multiplicity of external branches of given length in a random history of *n* leaves. A correspondence between the external branches of the ordered histories of size *n* and the non-peak entries of the permutations of size *n* −1 provides easy access to the length distributions of the first and second longest external branch in a random Yule history and coalescent tree of size *n*. The length of the longest external branch is also studied in dependence of root balance of a random tree. As a practical application, we compare the observed and expected number of mutations on the longest external branches in samples from natural populations.

## 1 Introduction

The Kingman coalescent is a fundamental model for population genetic analyses. In its original version [15, 16] it is the backward-in-time analogue of a pure birth process where each existing external branch is chosen uniformly to give rise to the next split into two offspring branches. As such, the involved trees are binary with internal nodes linearly ordered by time. Disregarding branch length and keeping track only of the ranking of the internal nodes, such trees are called (unlabeled) ranked trees [18] or histories [17], where the probability of a history of size *n* to be the underlying ranked tree topology of a random coalscent tree follows the Yule distribution [11, 22].

The stochastic, combinatoric, topological and population genetic properties of coalescent trees have been subject of numerous investigations. One prominent application in population genetics is to analyze and interpete the frequency spectrum of mutations in light of tree topology and of the length distribution of tree branches. In particular, singletons in the mutation frequency spectrum relate to the length of the external branches of the tree. Blum and François [3], as well as Caliebe *et al.* [5], have studied the length distribution of a randomly chosen external branch from a Kingman coalescent tree and derived also the limiting distribution for large *n*. The same topic has been investigated by Freund and Möhle [9] for the Bolthausen-Sznitman coalescent. These results have been generalized to the comprehensive class of Λ-coalescents by Diehl and Kersting [6], who also examined the asymptotic distribution of the external branch lengths ordered by size.

Here, we study the external branches of coalescent trees from a combinatorial point of view. We distinguish the time length of an external branch from its discrete length, the latter being defined as the rank (looking backward in time) of the parent node of the considered branch in the underlying history. An external branch of discrete length *s* is thus divided into *s* segments spanning the last *s* layers of the tree. When a random history of size *n* is selected under the Yule distribution, that is, it is the history underlying a random coalescent tree of *n* leaves, we derive several probabilistic properties of the length of its external branches. We focus on the probability of a given number of external branches of given length, on the probability that external branches of given length are absent, and on the probability of the length of the first and second longest external branches. Importantly, from the discrete length of an external branch, we can recover the probability density of its time length measured in coalescent units by summing exponentially distributed independent random variables.

Our study is also motivated by the practical question whether the observation of a certain number of singleton mutations in one single chromosome is compatible, or not, with the neutral infinite sites model [7, 14] of constant population size and constant mutation rate. We apply our results on the length of the two longest external branches of a tree to two kind of data: the mitochondrial genomes of three human populations [1], and to a nuclear gene of *Danio rerio* [20]. Non-recombining chromosomes, such as mitochondria, or short genomic fragments should not show any homogenizing effect, due to recombination, on the length distribution of external branches. Therefore, in the examples studied, we expect to recover and estimate the lengths of the longest and second-longest external branches of a single coalescent tree.

The paper is organized as follows. We introduce terminology and some useful properties of histories and coalescent trees in Section 2, showing in particular that external branch lengths in random trees can also be analyzed in terms of peaks of random permutations. In Section 3, we subdivide branches into branch segments. Given a random history of size *n*, i.e. with *n* leaves, we derive the counts—either 0, 1 or 2—of external branches with a given number of segments and ask in Section 4 how often a history misses external branches with a certain number of segments. We then consider the number of segments in the longest and second longest external branches (Section 5). Using convolution of exponential distributions, segment numbers can be scaled-back to coalescent time-units and the results be applied to experimental data (Section 6). We conclude with an outlook on some open problems (Section 7).

## 2 Histories, peaks of permutations and external branch length

Following [17], a *history* of size *n* is a full binary rooted tree with a ranking of its *n* − 1 internal nodes. If [*a, b*] denotes the set {*i* ∈ ℤ : *a* ≤ *i* ≤ *b*}, then the internal nodes of a history of size *n* are labeled by the integers in [1, *n* − 1], starting from the bottom (i.e., closest to the leaves) of the tree proceeding to the top (i.e., the root) in increasing order (Fig. 1). The root has thus label *n* − 1. A branch of a history is an edge connecting two internal nodes or one internal node and one leaf. In the latter case, the branch is said to be *external*. The ranking of the internal nodes of a history of size *n* divides the tree into *n* − 1 layers, with the *i*th layer intersecting exactly *i* branches. A branch *segment* is a part of a branch that crosses a given layer of the tree. For example, in the history depicted in Fig. 1 the two branches appended to the node labeled 6 consist of 1 and 4 branch segments. The external branch appended to node 3 consists instead of 3 branch segments. In general, the number of segments of an external branch of a history *t* is the label of the internal node of *t* from which the considered external branch descends.

**Figure 1:**
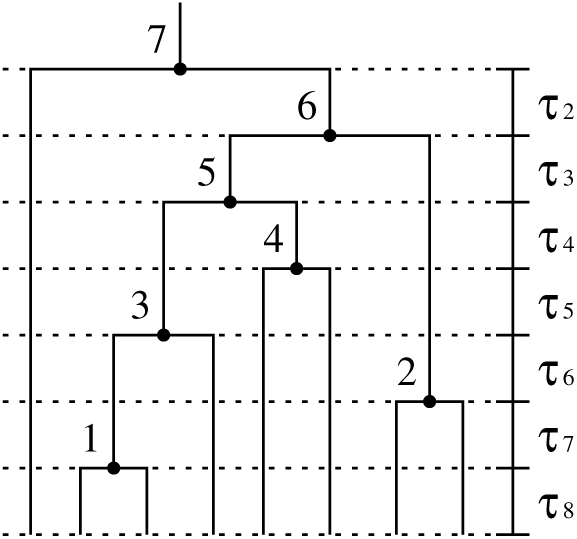
A history of size *n* = 8. Internal nodes are ranked and labeled by the integers in [1, *n* − 1], from bottom to top. The ranking divides the history into *n* − 1 layers, with the *i*th layer intersecting exactly *i* branches. A segment is a part of a branch that extends across a given layer. The number of segments of an external branch corresponds to the label of the internal node from which the branch descends. In a coalescent tree, an exponentially distributed variable *τ*_*i*_ assigns a time length to the *i*th layer of the history underlying the tree.

An *ordered* history of size *n* is a planar embedding of a history of size *n* in which subtrees have a left-right orientation. The *Yule* branching process [11, 22] creates a random ordered history of size *n* in *n* − 1 consecutive steps. Starting with a root branch, in the *i*th step of the process each one of the *i* present terminal nodes has the same probability to split into two new terminal nodes. After *n*−1 steps an ordered history is created with uniform probability among the (*n* − 1)! possible ordered histories of *n* leaves. By summing the probability 1*/*(*n* − 1)! of each ordered history with the same underlying (un-ordered) history, the uniform distribution over the set of ordered histories of size *n* induces a probability distribution—the Yule distribution—over the set of histories of size *n*. In particular, the Yule probability of a history *t* of size *n* can be seen as a function of the number |or(*t*)| of its different left-right orientations. More precisely, if |ch(*t*)| is the number of cherries (i.e., subtrees of size 2) in *t*, then |or(*t*)| = 2^*n*−1−|ch(*t*)|^—by flipping subtrees stemming from the *n* − 1 − |ch(*t*)| non-cherry internal nodes of *t*, we obtain the possible left-right orientations of *t*—and the Yule probability of the history *t* is given by |or(*t*)|*/*(*n* − 1)! [17]. In this manuscript, all random histories of fixed size *n* are selected under the Yule distribution, whereas ordered histories of size *n* are uniformly distributed.

The fact that the Yule distribution over the set of histories of size *n* is induced by the uniform distribution over the set of ordered histories of size *n* allows us to derive probabilistic properties of the number of branch segments present in the external branches of random histories by studying the number of branch segments in the external branches of random ordered histories of the same size. In particular, in the study of the number of segments of the external branches, combinatorial properties of ordered histories can be derived through a series of known enumerative results [4] on the number of permutations of fixed size with a given set of peak entries. Indeed, by bijectively encoding the (*n* − 1)! ordered histories of size *n* as permutations of *n* − 1 integers, the external branches of an ordered history *t* can be seen to correspond to the non-peak entries of the associated permutation *π*_*t*_, where the number of segments in each external branch of *t* is the value of the associated non-peak entry in the permutation *π*_*t*_. The mapping *t* → *π*_*t*_ is well known [10] and can be described as follows.

Given an ordered history *t* of size *n* with internal nodes labeled from bottom to top in increasing order (Fig. 1), let us consider an internal node of *t* labeled by the integer *k*. The permutation *π*_*t*_[*k*] associated with the subtree of *t* rooted at *k* is constructed recursively as *π*_*t*_[*k*] = (*π*_*t*_[*k*_*ℓ*_], *k, π*_*t*_[*k*_*r*_]), where *π*_*t*_[*k*_*ℓ*_], *π*_*t*_[*k*_*r*_] are the permutations associated with the subtrees of *t* rooted at the left and right children nodes *k*_*ℓ*_ and *k*_*r*_ of *k*, respectively. For example, if *t* is the ordered history of size *n* = 8 depicted in Fig. 1, then *π*_*t*_[7] = (7, 1, 3, 5, 4, 6, 2), where *π*_*t*_[7_*ℓ*_] = ∅ is the empty permutation and *π*_*t*_[7_*r*_] = *π*_*t*_[6] = (1, 3, 5, 4, 6, 2). Similarly, the permutation *π*_*t*_[6] has been constructed as *π*_*t*_[6] = (*π*_*t*_[6], 6, *π*_*t*_[6_*r*_)], where *π*_*t*_[6_*ℓ*_] = *π*_*t*_[5] = (1, 3, 5, 4) and *π*_*t*_[6_*r*_] = *π*_*t*_[2] = (2). In particular, the permutation *π*_*t*_ = *π*_*t*_[*n* − 1] of size *n* − 1 is by definition the permutation associated with the considered ordered history *t* of size *n*. Denoting by *π*_*t*_(*i*) the *i*th entry of the permutation *π*_*t*_, we say that *π*_*t*_(*i*) is a *peak* when *i* ≠ 1, *i* ≠ *n* − 1 and *π*_*t*_(*i* − 1) *< π*_*t*_(*i*) *> π*_*t*_(*i* + 1), and we observe the following property of the mapping *t* → *π*_*t*_: the entry *k* in the permutation *π*_*t*_ is a peak if and only if the node of *t* labeled by *k* has both its left and right child that are internal nodes of *t*. In other words, *k* is an *external* node of *t*, that is, *k* has *at least* one descending external branch, if and only if the entry *k* in the permutation *π*_*t*_ is not a peak. For instance, the peak entries of the permutation *π*_*t*_ = (7, 1, 3, 5, 4, 6, 2) associated with the ordered history *t* depicted in Fig. 1 are 5 and 6, which correspond to the internal nodes of *t* without a descending external branch.

The study of probabilistic properties of the number of segments of the external branches of a random history can assist in the analysis of the time length of the external branches of a Kingman coalescent tree [12, 15, 21]. A coalescent tree of size *n* can be modeled as a random history *t* of *n* leaves and a sequence (*τ*_2_, …, *τ*_*n*_) of independent exponentially distributed random variables assigning a time length to the different layers of *t* (Fig. 1) [19, 23]. Under the model, the variable *τ*_*i*_ has mean 𝔼 [*τ*_*i*_] = 1*/λ*_*i*_, with 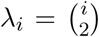. From 𝔼[*τ*_*i*_] we can easily recover the expected value of the time length *τ* (*s*) of an external branch of *t* containing exactly *s* segments. The expectation of *τ* (*s*) is the sum of the expectations of the time length of the last *s* layers of the history *t*, that is,

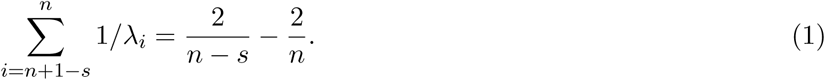

More generally, the probability density function *f*_*s*_(*x*) of the time length *τ* (*s*) is the density of the sum 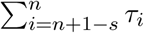, which is given [2, 8] by

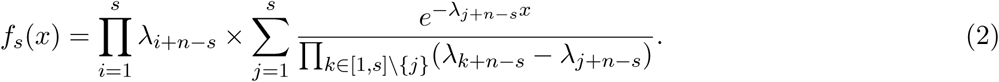

The latter formula enables the calculation of the probability of the time length of an external branch of a coalescent tree given the number of segments possessed by that branch in the underlying history.

## 3 The number of external branches with a given number of segments

In this section, we study the number of ordered histories of size *n* with *µ* ∈ {0, 1, 2} external branches consisting of exactly *s* segments. In particular, dividing this number by (*n* − 1)!—i.e., by the total number of ordered histories of *n* taxa—we obtain the probability that a random history of size *n* selected under the Yule distribution has *µ*_*s*_ external branches of *s* segments. This calculation is then extended to the conditional probability that a Yule-distributed history of size *n* has *µ* external branches of *s* segments given that it has *µ*_*r*_ external branches of *r* segments.

Let *a*_*n,s,µ*_ denote the number of ordered histories of size *n* with *exactly µ* external branches of 1 ≤ *s* ≤ *n* − 1 segments. Constructing ordered histories of size *n* ≥ 3 by splitting a leaf of an ordered history of size *n* − 1 yields for 1 *< s* ≤ *n –* 1

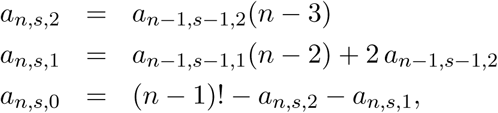

where *a*_*n*,1,2_ = (*n* − 1)! and *a*_*n*,1,1_ = 0 for every *n* ≥ 2. For example, the recurrence for *a*_*n,s*,1_ generates a tree of size *n* with exactly one external branch of *s* segments either from a tree of size *n* − 1 with exactly one external branch of *s* − 1 segments by splitting one of its *n* − 2 external branches of length different from *s* − 1 or from a tree of size *n* − 1 with 2 external branches of *s* − 1 segments by splitting one of these two branches. Note, that when we split a branch, all the remaining branches increase their number of segments by one.

Solving the recurrences above we find

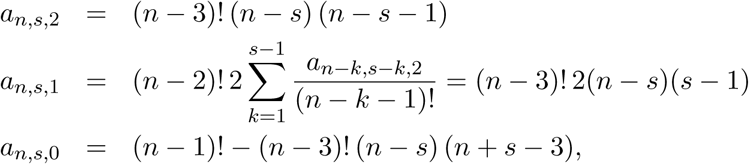

and the probability *p*_*s*_(*µ*) of a history of size *n* with *µ* external branches of *s* segments is given by

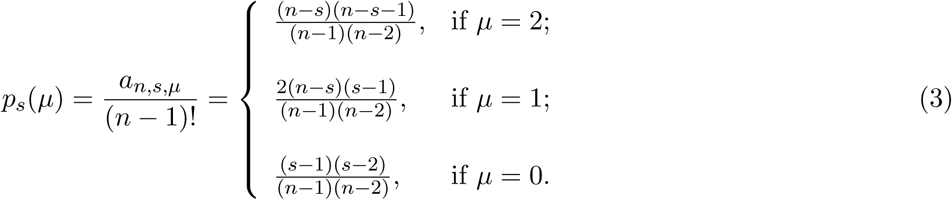

As shown in Fig. 2, we have the symmetries *p*_*s*_(0) = *p*_*n*−*s*+1_(2), *p*_*s*_(1) = *p*_*n*−*s*+1_(1).

**Figure 2:**
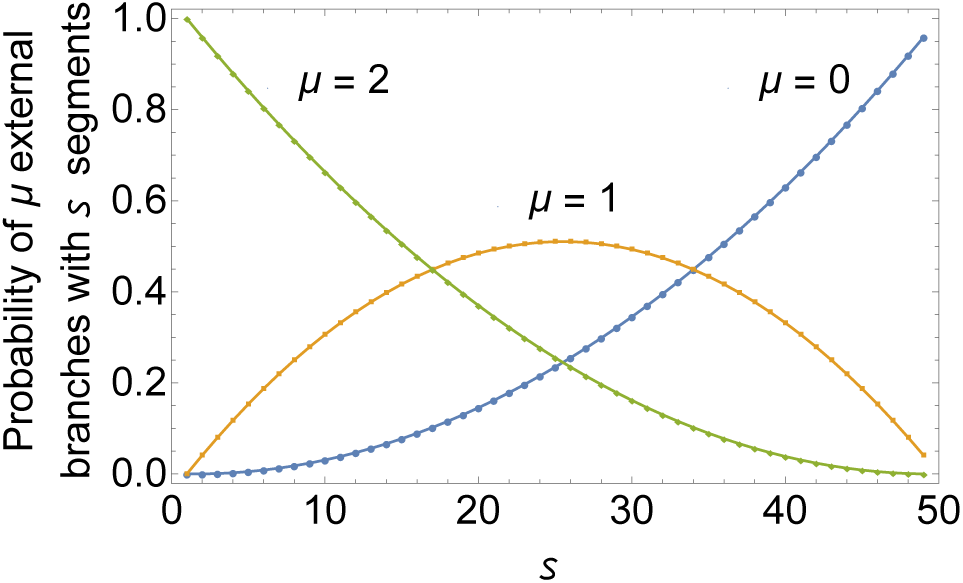
Probability that a random Yule-distributed history of size *n* = 50 has *µ* ∈ {0, 1, 2} external branches with *s* ∈ [1, *n* − 1] segments. The probability increases (resp. decreases) with *s*, when *µ* = 0 (resp. *µ* = 2).

The expected number of external branches with *s* segments in a random history of size *n* decreases linearly with *s* as given by 𝔼 _*s*_[*µ*] = 2(*n* − *s*)*/*(*n* − 1). Furthermore, we can calculate the value *s*^*^ = *s*^*^(*n*) such that *p*_*s*_(0) ≤ 1*/*2 for every *s* ≤ *s*^*^. We find

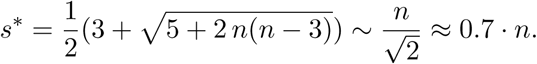

In other words, for a random history of size *n*, the probability of having at least one external branch with *s* segments is larger or smaller than 50% depending on whether *s < s*^*^ or *s > s*^*^, respectively. Because *p*_*s*_(0) = *p*_*n*−*s*+1_(2), we also have that *p*_*s*_(2) is larger or smaller than 50% when *s < n* − *s*^*^ + 1 or *s > n* − *s*^*^ + 1, respectively, where 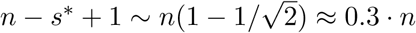.

### Conditional probability calculation

The calculation above can be extended to the number 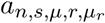 of ordered histories of size *n* in which there are *µ* ∈ {0, 1, 2} external branches of *s* segments and *µ*_*r*_ *∈* {0, 1, 2} external branches of *r* segments. When 1 *< s, r* ≤ *n* − 1, we have the following recurrences

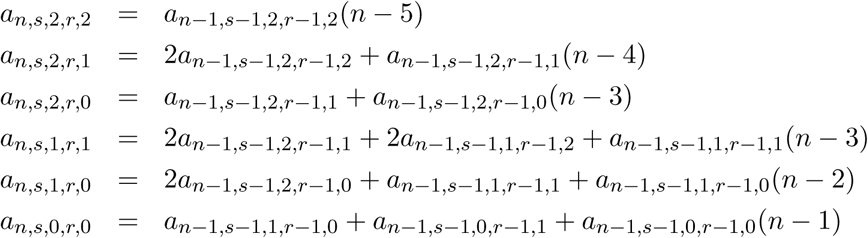

where 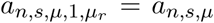 if 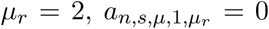 if 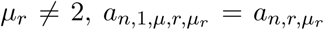 if *µ* = 2, and 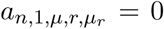 if *µ* ≠ 2. For instance, the recurrence for *a*_*n,s*,1,*r*,1_ yields a tree of size *n* with exactly one external branch of *s* segments and exactly one external branch of *r* segments either from a tree of size *n* − 1 with exactly one external branch of *s* − 1 segments and one external branch of *r* − 1 segments by splitting one of its *n* − 3 external branches of length different from *s* − 1 and *r* − 1 or from a tree of size *n* − 1 with 2 external branches of *s* − 1 (or *r* − 1) segments and one external branch of *r* − 1 (or *s* − 1) segments by splitting one of the two branches with *s* − 1 (or *r* − 1) segments.

Solving the recurrences yields for *n* ≥ 5 and 1 ≤ *s, r* ≤ *n* − 1 the following formulas

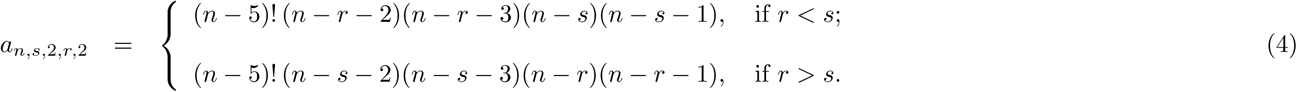

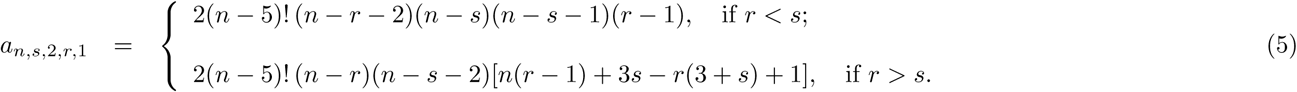

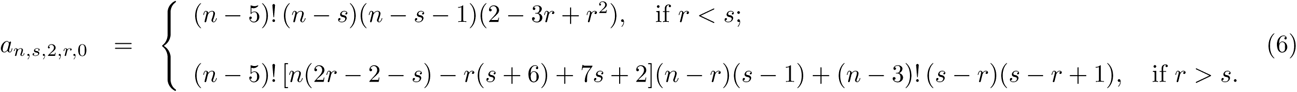

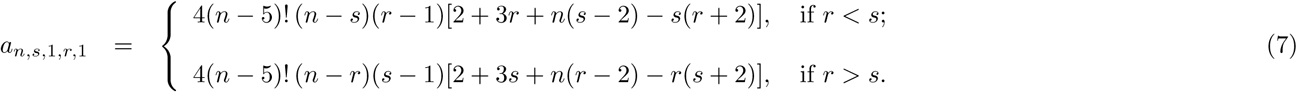

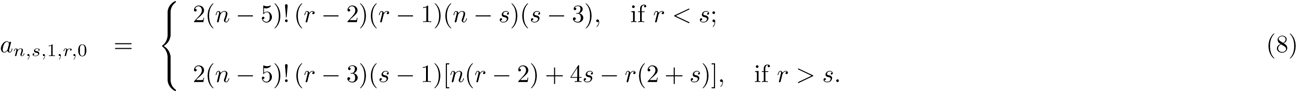

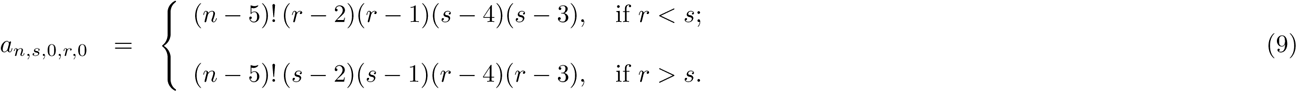

Therefore, for a random history with *n* ≥ 5 taxa, the conditional probability *p*_*s*_(*µ*|*r, µ*_*r*_) of *µ* ∈ {0, 1, 2} external branches of *s* segments given *µ*_*r*_ *∈* {0, 1, 2} external branches of *r* segments can be computed as

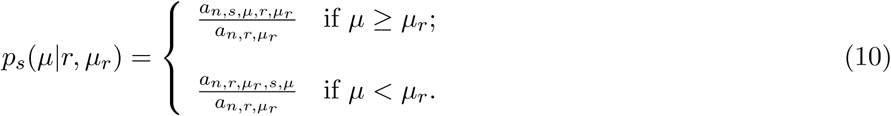

In Fig. 3, we plot *p*_*s*_(*µ*|*r, µ*_*r*_) (solid line) and *p*_*s*_(*µ*) (boxes) for *n* = 50. When *r* = 10 and *µ*_*r*_ = 0 (left column), we see that *p*_*s*_(0|*r, µ*_*r*_) ≤ *p*_*s*_(0), *p*_*s*_(1|*r, µ*_*r*_) ≤ *p*_*s*_(1) if *s* ≤ *n/*2, and *p*_*s*_(2|*r, µ*_*r*_) ≥ *p*_*s*_(2). Thus, a random history that misses a short (*r* = 10) external branch, has a slightly smaller probability to miss an external branch of *s* segments than a random unconstrained history of the same size. Interestingly, the missing short external branch of *r* segments does not increase, for *s* close to *r*, the probability of having one external branch of *s* segments—in fact, *p*_*s*_(1|*r, µ*_*r*_) *< p*_*s*_(1)—but it increases the probability of having two external branches of *s* segments. When instead *r* = 40 and *µ*_*r*_ = 2 (right column of Fig. 3), we see that the existence of two long (*r* = 40) external branches decreases, for *s* close to *r*, both the probability of having two external branches of *s* segments and the probability of having one external branch of *s* segments.

**Figure 3:**
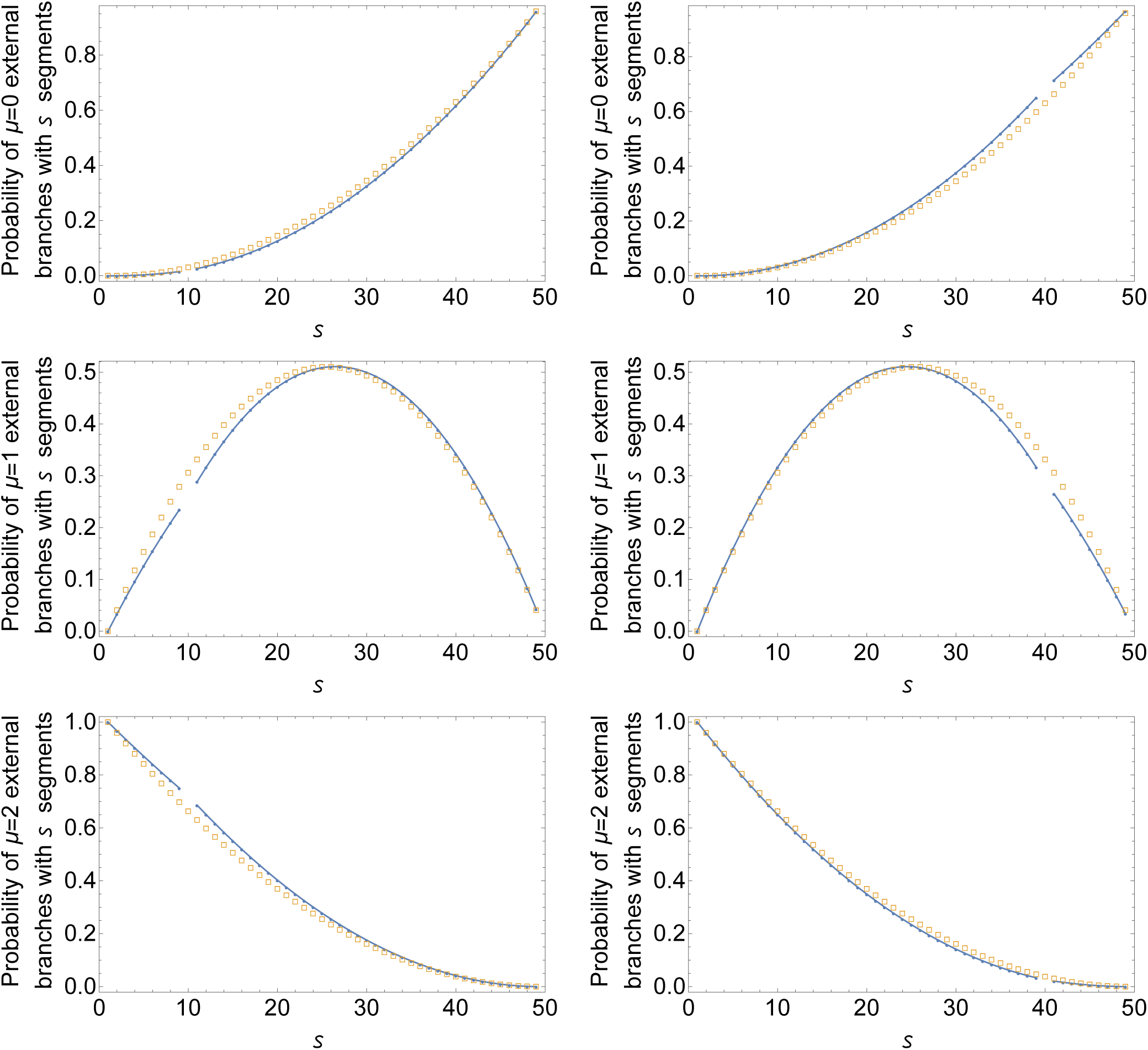
Conditional probability *p*_*s*_(*µ*|*r, µ*_*r*_) of a random Yule-distributed history of size *n* = 50 having *µ* ∈ {0, 1, 2} external branches of *s* ∈ [1, *n* − 1] segments. In the left column, *r* = 10 and *µ*_*r*_ = 0. In the right column, *r* = 40 and *µ*_*r*_ = 2. Boxes give the probability *p*_*s*_(*µ*) in an unconstrained history.

## 4 The probability that only the external branches of length *s* are absent

Here, we provide a procedure for evaluating the probability that a random history of size *n* does not have external branches of *s* segments if and only if *s* belongs to a given integer set. Results of this section are derived by considering ordered histories and the correspondence between their external branches and the non-peak entries of the associated permutations (Section 2).

### Histories with few missing external branches

We recall from Section 2 that a node of a history is said to be *external* when it has at least one child that is a leaf. The external branches of a history are therefore those branches that are appended too an external node. In particular, when we label the nodes of a history of size *n* from 1 to *n* − 1 moving upwards in the tree (Fig. 1), the number of segments of an external branch is given by the integer label of the external node to which the branch is appended. For a given history *t* with ⌈*n*/2⌉ ≤ *k* ≤ *n* − 1 external nodes, let *ε* _*t*_ ≡ {*e*_1_, …, *e*_*k*_} be the set of labels of the external nodes of *t*. Thus, *t* has at least one external branch with *s* segments if and only if *s* ∈ *ε* _*t*_, and we say that *t* misses an external branch of *s* segments when *s* ∉ *ε*_*t*_.

When *ε* ⊆ [1, *n* − 1] is a set of *k* ∈ {*n* − 1, *n* − 2, *n* − 3} positive integers, the probability that a random history of size *n* has its set of missing external branches given by 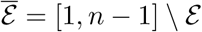 can be calculated as

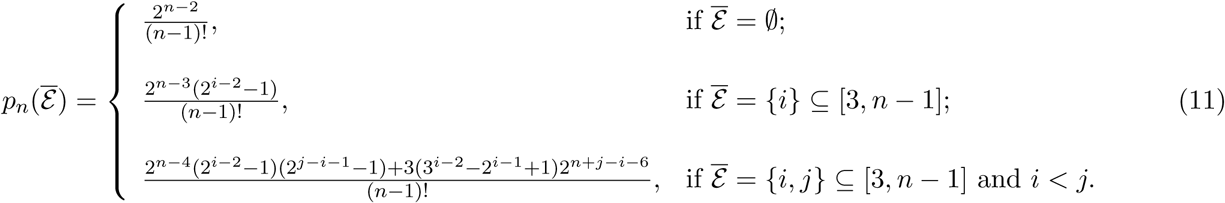

In particular, as shown in Theorem 3.1 of [4], the enumerator in each formula counts the number of permutations of size *n* − 1—i.e., the ordered histories of size *n*—in which the peak entries—i.e., the non-external nodes— are exactly those belonging to the set 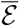. We remark that the probability 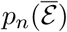 is different from the type of probabilities analyzed in Section 3. For instance, when 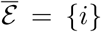 the probability 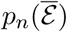 in (11) considers the histories of size *n* in which the only non-external node is *i*. The probability *p*_*i*_(0) of Eq. (3) is instead the probability of an history in which at least the node *i* is not external—or, equivalently, in which there are 0 external branches of *i* segments.

### Existence and probability of a history with a given set of missing external branches

We first give a necessary and sufficient condition for the existence of at least one history of size *n* with a given set of missing external branches. If *ε* ⊆ [1, *n* − 1] and 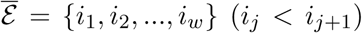 (*i*_*j*_ *< i*_*j*+1_), then there exists at least one history *t* of size *n* ≥ 4 such that *ε*_*t*_ = *ε* if and only if either *w* = 0—i.e. there are no missing external branches—or *i*_*j*_ ≥ 2*j* + 1 for every 1 ≤ *j* ≤ *w*—i.e. the number of segments of the *j*th smallest missing external branch is larger than or equal to 2*j* + 1. This characterization is given in Theorem 2.1 of [4] in terms of allowed peak sets of a permutation of given size.

When *n* ≥ 4 and 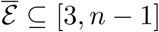 satisfies the condition above, the probability 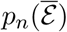 that a random history of size *n* has its set of missing external branches given by 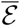 can be calculated recursively as follows

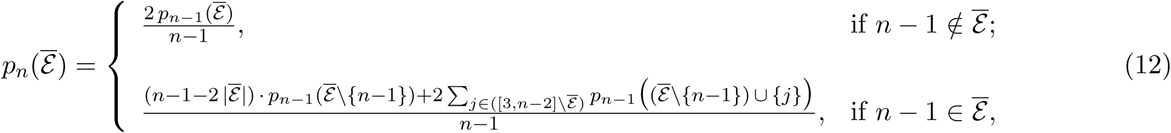

where *p*_4_(∅) = 2*/*3, *p*_4_({3}) = 1*/*3, and 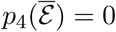 otherwise. The second formula in (12) follows from Lemma 3.2 of [4]. The first formula is instead a direct consequence of the fact that an ordered history of size *n* having an external branch of *n* − 1 segments and a set 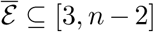 of missing external branches (Fig. 4) must have as left or right root subtree an ordered history of size *n* − 1 whose set of missing external branches is 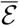. In Fig. 4 (right), we plot for *n* = 8 the probability 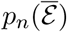 for all the 20 admissibe sets 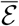 of missing external branches. Among sets 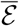 of the same cardinality, we observe a correlation between the probability of each 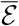 and its ranking in the lexicographic order. This correlation is weaker if we compare the probabilities of sets 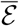 with a different number of elements. For example, {5, 7} is lexicographically smaller than {6} but it has a larger probability. Similarly, the probability of {6, 7} is larger than the probability of {7}.

**Figure 4:**
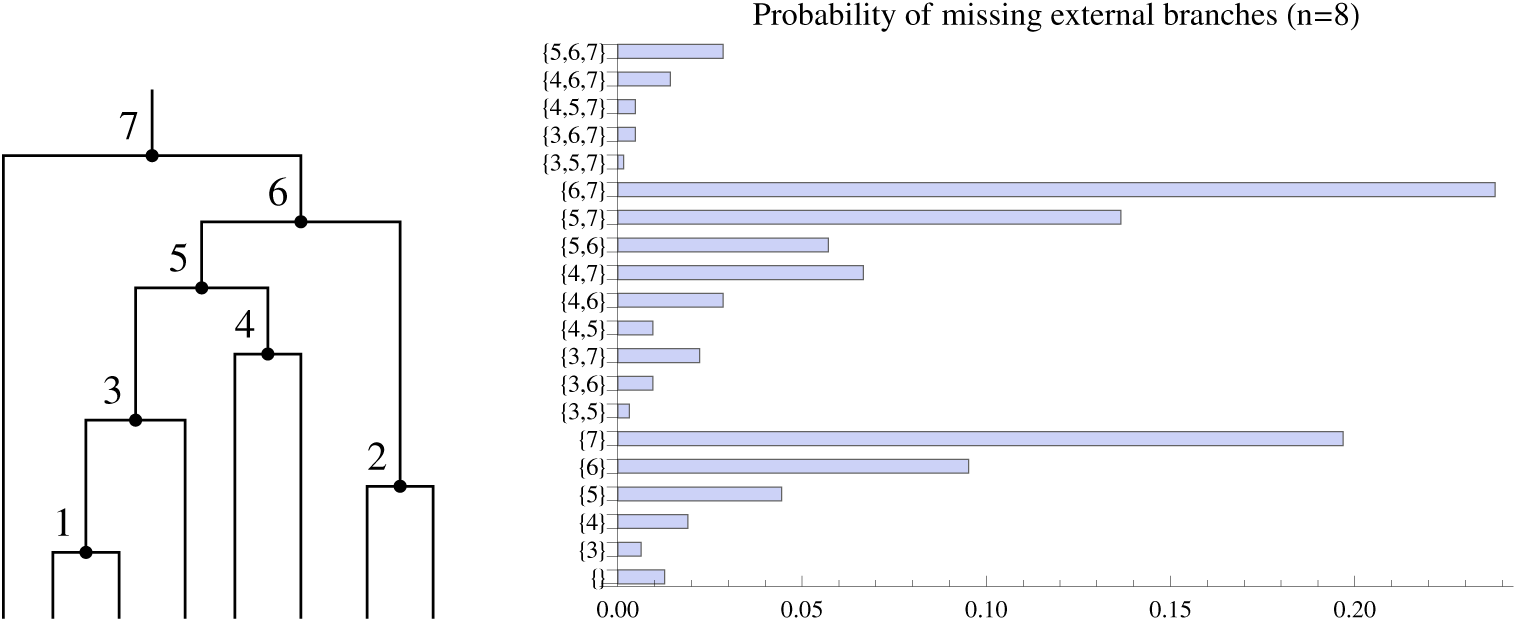
Left: a history of size *n* = 8 whose set of missing external branches is 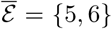. In the tree, there is at least one external branch of *s* segments if and only if 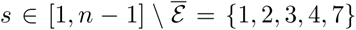. Right: probability of all the possible admissible sets of missing external branches for a random Yule-distributed history of size *n* = 8.

The recursive procedure described in (12) can be used to extend the calculation performed in Section 3 of the joint probability *a*_*n,s*,0,*r*,0_*/*(*n* − 1)! of missing two external branches of *s* and *r* segments in a random history of size *n*. More precisely, given a set *S* ⊆ [3, *n* − 1], the probability that a random history of size *n* misses *at least* those branches with discrete length listed in *S* can be evaluated as 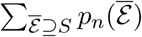, that is, by summing the probabilities of the admissible sets 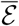 containing *S*.

## 5 The number of segments in the first and second longest external branches

In this section, we study the random variables *s′* and *s″* counting, respectively, the number of branch segments in the longest and second longest external branch of a random history of given size. In Section 5.1, we provide an exact formula for the probability of a given value of *s′* ∈ [⌈*n/*2⌉, *n* − 1]. The probability of *s′* to be smaller than a certain threshold is studied in Section 5.1.1. Relationships between *s′* and tree imbalance are analyzed in Section 5.1.2. In Section 5.2, we calculate the probability of a given value of *s″* and the conditional probability of *s″* given *s′* for a random history of fixed size.

### 5.1 The number of segments in the longest external branch

In this section, we calculate the probability that the longest external branch in a random history of given size has *s′* = *s* segments. As in the previous sections, we use the equivalence with the uniform distribution over ordered histories of size *n* or permutations of size *n* − 1.

Let Π_*n*_(*X*) denote the number of permutations of size *n* with peak entries matching the elements of the set *X* ⊆ [3, *n*], and choose *s* ∈ [*I*(*n* + 1)*/*2*1, n*]. For a fixed set *S* ⊆ [3, *s* − 1], by setting *k* = *n* − *s* in Lemma 3.3 of [4], for *n* ≥ 2 we find

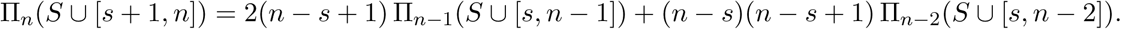

The latter equation relates the number of permutations of size *n* with peak entries given by the elements of the set *S* ∪ [*s* + 1, *n*] to the number of permutations of size *n* − 1 and *n* − 2 with peak entries given by *S* ∪ [*s, n* − 1] and *S* ∪ [*s, n* − 2], respectively. Note that if *s* = *n* and *n* − 1 ∈ *S*, then Π_*n*−2_(*S* ∪ [*s, n* − 2]) = 0.

If we sum both sides of the latter equation over the possible subsets *S* of [3, *s* − 1], then we obtain

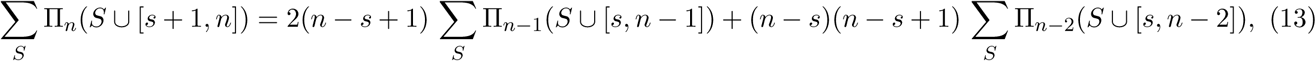

where the sum Σ_*S*_ Π_*n*_(*S* ∪ [*s* + 1, *n*]) counts the permutations of size *n* in which the largest non-peak entry is *s*, and the sums Σ_*S*_ Π_*n*−1_(*S* ∪ [*s, n* − 1]) and Σ_*S*_ Π_*n*−2_(*S* ∪ [*s, n* − 2]) count respectively the permutations of size *n* − 1 and *n* − 2 in which the largest non-peak entry is strictly smaller than *s*. For instance, set *n* = 5. For *s* = 3, Table 2 of [4]—which reports the number of permutations of size *n* ≤ 8 with a fixed set of peak entries—gives Σ_*S*_ Π_*n*_(*S* ∪ [*s* +1, *n*]) = 12, Σ_*S*_ Π_*n*−1_(*S* ∪ [*s, n* − 1]) = 0 and Σ_*S*_ Π_*n*−2_(*S* ∪ [*s, n* −2]) = 2, where 12 = 2 · 3 · 0+2 · 3 · 2 in agreement with Eq. (13). For *s* = 4, we have Σ_*S*_ Π_*n*_(*S* ∪ [*s* + 1, *n*]) = 60, Σ_*S*_ Π_*n*−1_(*S* ∪ [*s, n* − 1]) = 12 and Σ_*S*_ Π_*n*−2_(*S* ∪ [*s, n* − 2]) = 6, where 60 = 2 · 2 · 12 + 1 · 2 *·* 6. Finally, for *s* = 5 we have Σ_*S*_ Π_*n*_(*S* ∪ [*s* + 1, *n*]) = 48, Σ_*S*_ Π_*n*−1_(*S* ∪ [*s, n* − 1]) = 24 and Σ_*S*_ Π_*n*−2_(*S* ∪ [*s, n* − 2]) = 6, where 48 = 2 · 1 · 24 + 0 · 1 · 6.

Note that the number Σ_*S*_ Π_*n*_(*S* ∪ [*s* + 1, *n*]) of permutations of size *n* in which the largest non-peak entry is *s* can be seen as the difference between the number of permutations of size *n* whose largest non-peak entry is strictly smaller than *s* + 1 and the number of permutations of size *n* whose largest non-peak entry is strictly smaller than *s*. Hence, by using the correspondence between non-peak entries of permutations of size *n* and external branches of ordered histories of size *n* + 1, from (13) we have the following equation for the number *a*_*n,s*_ of ordered histories of size *n* in which the longest external branch has a number of segments *strictly* smaller than *s*:

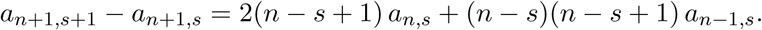

Replacing *n* + 1 by *n* and *s* + 1 by *s* in the latter equation, for *n* ≥ 3 we obtain the recurrence

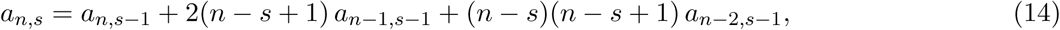

where *a*_*n,s*_ = 0 if *s* ≤ ⌈*n/*2⌉, and *a*_*n,s*_ = (*n* − 1)! if *s* = *n*. By iteratively setting *s* = *n, s* = *n* − 1, *s* = *n* − 2, … in Eq. (14) and extracting the term *a*_*n,s*−1_, we can recursively calculate a formula for *a*_*n,n*−*i*_. For the first values of *i* ∈ [0, 5], we find

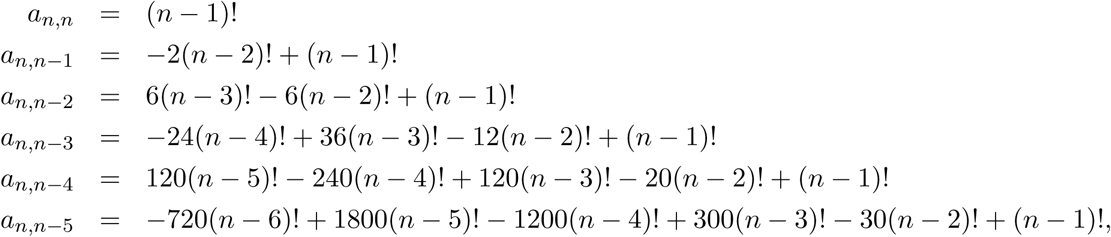

and more in general, as shown in the Appendix, we have

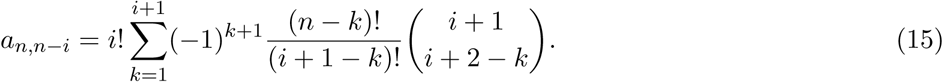

Setting *s* = *n* − *i*, the latter formula can be rewritten as

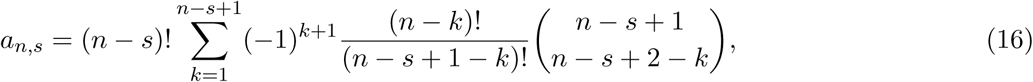

and for *n* ≥ 3 the probability *p*_*n*_(*s*) that a random history of size *n* has its longest external branch containing exactly *s′* = *s* segments (Fig. 5, left) can be computed as

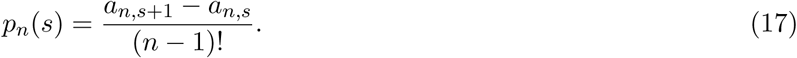

**Figure 5:**
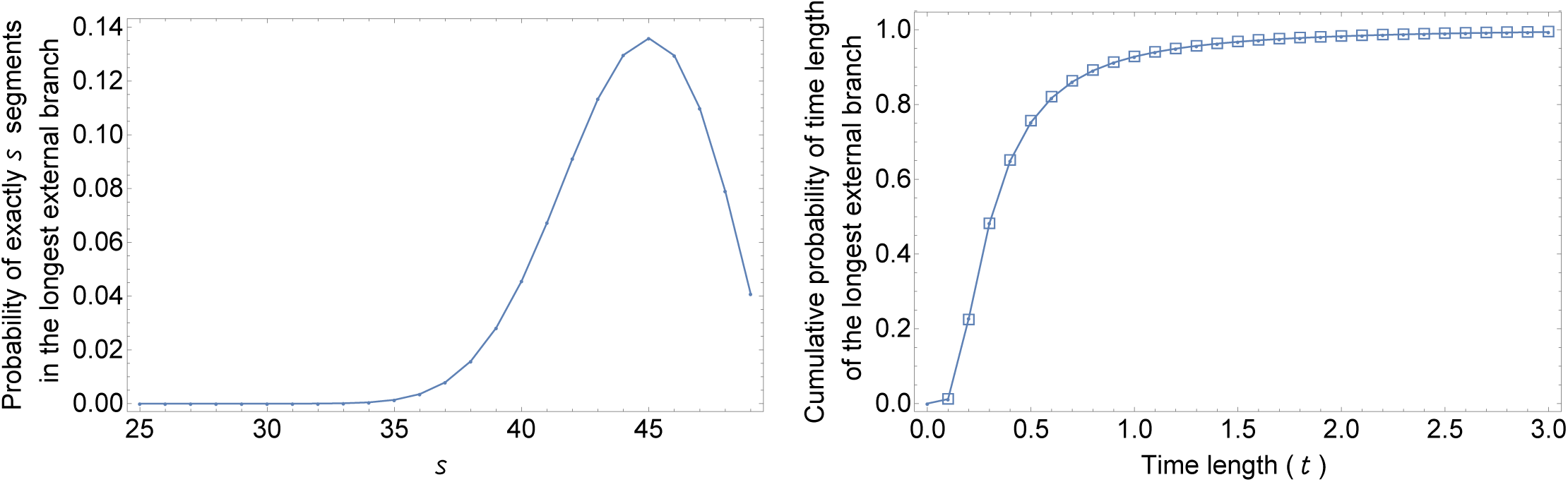
Left: Probability that a Yule-distributed history of size *n* = 50 has exactly *s* ∈ [⌈ *n/*2 ⌉, *n* − 1] segments in its longest external branch. Right: Cumulative probability that the longest external branch of a history of size *n* = 50 has length at most *t* (in coalescent time units). Theoretical probabilities (solid line) are calculated by Eq. (17), using Eq. (2) with 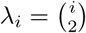. Simulated data (boxes) have been obtained through `ms` [13].

By using Eqs. (2) and (17), the probability that the time length *τ* of the longest external branch in random history of size *n* lies in the interval *a < τ* ≤ *b* can be calculated as

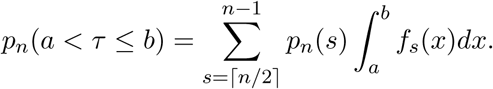

In Fig. 5 (right), the cumulative probability *p*_*n*_(*τ* ≤ *t*) is plotted setting *n* = 50 and letting *t* range over the interval [0, 3] in steps of 0.1. The theoretical line is in perfect agreement with rescaled data obtained through the `ms` coalescent simulator [13]. Note that in the ms setting, the mean length of the *i*th layer of a coalescent tree is 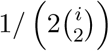, while in our theoretical calculations the mean is 1*/λ*_*i*_, with 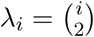.

#### 5.1.1 The longest external branch has a large number of segments

The probability that the longest external branch of a random Yule-distributed history of size *n* has exactly *s* segments has been calculated in the previous section. The plot given in Fig. 5 (left) shows that for a large fraction of the ordered histories of size *n* the longest external branch has a number of segments quite close to the maximum value *n* − 1. To better understand this observation, we derive in this section an approximation for the value *d*_*α*_ such that with probability *α* a random history of size *n* has less than *n* − *d*_*α*_ segments in its longest external branch.

From the left-top plot of Fig. 3, we see that missing an external branch of size *r* has only a small effect on the probability of missing an external branch of size *s* ≠ *r*. For a given value of *d* ≥ 1, we approximate the probability Prob(*µ*_*n*−1_ = … = *µ*_*n*−*d*_ = 0) that for all *i* ∈ [1, *d*] a random history of size *n* has *µ*_*n*−*i*_ = 0 external branches of length (number of segments) *n* − *i* as the product 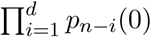 of the probabilities *p*_*n*−*i*_(0) given in Eq. (3). That is,

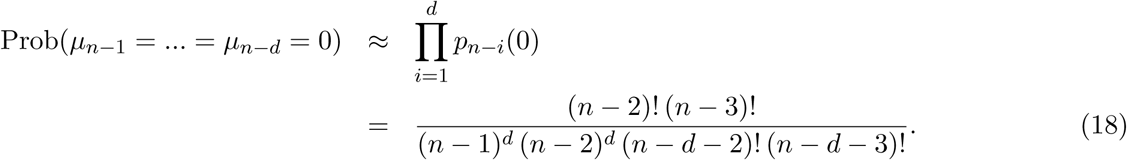

Note that Prob(*µ*_*n*−1_ = … = *µ*_*n*−*d*_ = 0) can be seen as the probability of a random history of size *n* with its longest external branch containing less than *n* − *d* segments, and its exact value can be calculated from Eq. (16) as Prob(*µ*_*n*−1_ = … = *µ*_*n*−*d*_ = 0) = *a*_*n,n*−*d*_*/*(*n* − 1)!. The accuracy of the approximation in (18) can be verified for *n* = 100 in Fig. 6 (left). The figure also shows that for a fixed value 0 ≤ *α* ≤ 1, the probability Prob(*µ*_*n*−1_ = … = *µ*_*n*−*d*_ = 0) is equal to *α* for a value of *d* much smaller than *n*. For instance, if we set *α* = 0.2 with *n* = 100, then Prob(*µ*_*n*−1_ = … = *µ*_*n*−*d*_ = 0) = *α* for *d* ≈ 12. In other words, there is an 80% probability that a random Yule-distributed history of size *n* = 100 has at least 100 − 12 = 88 segments in its longest external branch. To measure the probability that the longest external branch of a random history is shorter than a certain threshold, we use the approximation in Eq. (18) for studying the value *d*_*α*_ of *d* such that

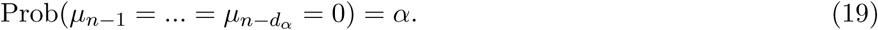

**Figure 6:**
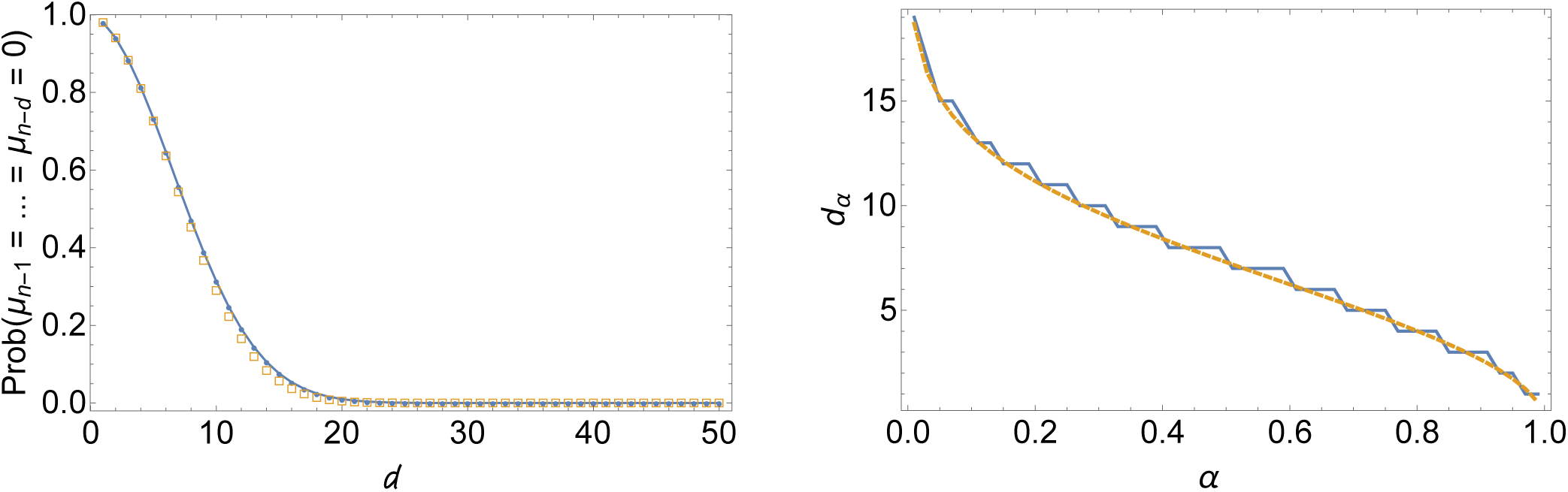
Left: The probability of 0 external branches of size larger than or equal to *n* − *d*—or the probability of the longest external branch with less than *n* − *d* segments—in a random Yule-distributed history of size *n* = 100. Boxes give the exact probability evaluated as *a*_*n,n*−*d*_*/*(*n* − 1)! by using Eq. (16), the solid line is the approximation given in Eq. (18). Right: Plot of *d*_*α*_ for *n* = 100 and 0.01 ≤ *α* ≤ 0.99 (in steps of 0.02). The zigzag line is *d*_*α*_ computed as the integer *d* for which the probability Prob(*µ*_*n*−_1 = … = *µ*_*n*−*d*_ = 0) is closest (not necessarily equal) to *α*. The smooth line is the approximation of *d*_*α*_ given in Eq. (21).

We find that *d*_*α*_ grows roughly like a constant multiple of 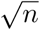, with the constant depending on the chosen *α*.

Assuming *d*_*α*_*/n* ≈ 0 for a fixed *α* and *n* sufficiently large, we apply Stirling’s approximation 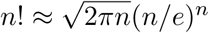 to Eq. (18) and, from (19), we obtain

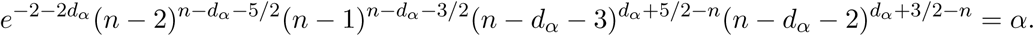

Taking the logarithm of the latter expression gives the equation

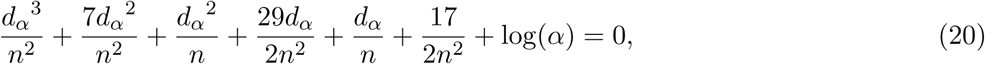

where we have used the second order approximations log(*n* − 2) ≈ log(*n*) − 2*/n* − (−2*/n*)^2^*/*2, log(*n* − 1) ≈ log(*n*) − 1*/n* − (−1*/n*)^2^*/*2, log(*n* − *d*_*α*_ − 3) ≈ log(*n*) − (*d*_*α*_ + 3)*/n* − [−(*d*_*α*_ + 3)*/n*]^2^*/*2, log(*n* − *d*_*α*_ − 2) ≈ log(*n*) − (*d*_*α*_ + 2)*/n* − [−(*d*_*α*_ + 2)*/n*]^2^*/*2. Note that −*d*_*α*_^3^*/n*^2^ − *d*_*α*_^2^*/n* = −*d*_*α*_^2^*/n*(*d*_*α*_*/n* + 1) ≈ −*d*_*α*_^2^*/n* is the leading term in the left-hand side of Eq. (20), where all the remaining terms of that side of the equation are close to 0 if *d*_*α*_*/n* ≈ 0. Thus, in order to satisfy the latter equation, −*d*_*α*_^2^*/n* must be close to log(*α*), that is, 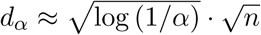 for *n* large. Approximating *d*_*α*_ as a sum of powers of 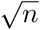 we find

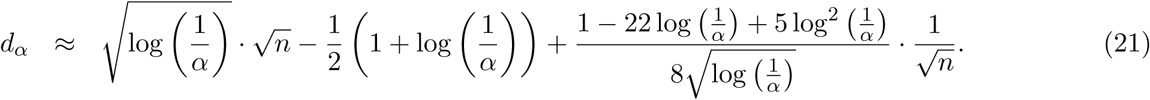

In particular, the estimate in (21)—plotted in Fig. 6 (right) for *n* = 100 and different values of *α*—has been obtained by first substituting 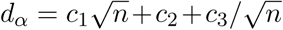 in the left-hand side of the equation given in (20), and then requiring the first three largest terms of the resulting expression—i.e., the coefficients of *n*^−*i/*2^ for *i* = 0, 1, 2—to be identically 0. The values of *c*_1_, *c*_2_, and *c*_3_ found in this way are such that the left-hand side of the polynomial equation in (20) is equal to 0 up to an error term of order *O*(1*/n*^3*/*2^). Higher precision can be obtained by the substitution 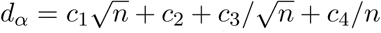, yielding 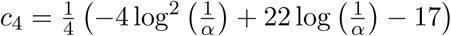 with an error term in the equation of order *O*(1*/n*^2^).

The square root behavior of the quantity *d*_*α*_(*n*) shows that for increasing tree size a random history will have with high probability a large number of segments in its longest external branch. For instance, setting *α* = 0.2 in (21) we obtain the estimate 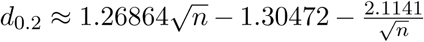, where the longest external branch of a random history of size *n* has at least *n* − *d*_0.2_ segments with probability 1 − *α* = 0.8.

#### 5.1.2 Longest external branch and root imbalance

In this section, we study how root imbalance affects the number of segments of the longest external branch. Our calculations show that the length of the longest external branch is almost independent of imbalance and affected only by extreme values of the latter parameter. In order to measure root imbalance of a history *t* of size *n*, we consider the parameter *ω*(*t*) *∈* [1, ⌋*n/*2⌋] defined as the size of the smallest root subtree of *t*. For instance, if *t* is the history of Fig. 1, then *ω*(*t*) = 1. Also in this section, we use the fact that the Yule distribution over histories of *n* leaves is induced by the uniform distribution over ordered histories of *n* leaves (Section 2).

If *t* is an ordered history of size *n*, let *t*_1_ and *t*_2_ be the *rescaled* left and right root subtrees of *t* of size *n*_1_ and *n*_2_, respectively. If the left root subtree *t*_*ℓ*_ of *t* has size *n*_1_, then *t*_1_ is obtained from *t*_*ℓ*_ by relabeling its *n*_1_ − 1 internal nodes with the integers in [1, *n*_1_ − 1]. Each internal node receives the new label *i* ∈ [1, *n*_1_ − 1] if the same node has the *i*th largest label when considered in *t*_*ℓ*_. Similarly, for the rescaled right root subtree *t*_2_ of *t*. As an example, consider the ordered history *t* given by the right root subtree of the history depicted in Fig. 1. In Newick format, *t* = ((((•, •)_1_, •)_3_, (•, •)_4_)_5_, (•, •)_2_)_6_, where the integer next to a closed parenthesis is the label of the associated internal node. The left root subtree of *t* is given by *t*_*R*_ = (((•, •)_1_, •)_3_, (•, •)_4_)_5_. The rescaled left root subtree is *t*_1_ = (((•, •)_1_, •)_2_, (•, •)_3_)_4_, which is obtained by replacing the labels 3, 4, 5 by 2, 3, 4, respectively, in *t*_*R*_. The rescaled right root subtree of *t* is instead *t*_2_ = (•, •)_1_, which is obtained by taking the right root subtree (•, •)_2_ of *t* with the label 2 replaced by 1. Finally, let 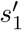 (resp. 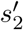) be the number of segments in the longest external branch of *t*_1_ (resp. *t*_2_), and denote by 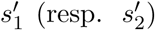 the number of segments in the longest external branch of the non-rescaled left (resp. right) root subtree of *t*. Thus, 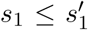 and 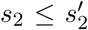. For example, if *t* is the history given by the right root subtree of the history depicted in Fig. 1, then we have 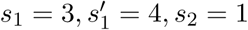, and 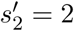.

Let 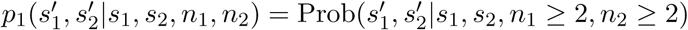. This is the joint probability of a given number of segments in the longest external branch of the non-rescaled left and right root subtree of a random ordered history of size *n* = *n*_1_ + *n*_2_, given the number of segments *s*_1_ and *s*_2_ in the longest external branch of its rescaled left and right root subtree. The tree decomposition depicted in Fig. 7 yields

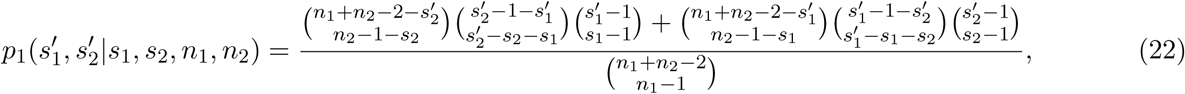

where we set 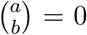 if *a <* 0 or *b <* 0. In particular, given two ordered histories *t*_1_ and *t*_2_ of size *n*_1_ ≥ 2 and *n*_2_ ≥ 2, respectively, with *s*_1_ and *s*_2_ segments in their longest external branch, we consider the ordered histories *t* of size *n* = *n*_1_ + *n*_2_ having *t*_1_ and *t*_2_ as rescaled root subtrees. The number of such histories *t* is given by the denominator of the latter formula, which counts the possible orderings of the *n*_1_ − 1 internal nodes of *t*_1_ with the *n*_2_ − 1 internal nodes of *t*_2_. The enumerator of the same formula counts the orderings of the internal nodes of *t*_1_ and *t*_2_ for which the resulting history *t* is compatible with the values of 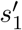 and 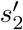. More precisely, when the node *s*_1_ of *t*_1_ has a lower rank in *t* than the rank assigned to the node *s*_2_ of *t*_2_ (Fig. 7A), the number of orderings is given by the first summand in the enumerator of Eq. (22). When instead *s*_1_ is placed above *s*_2_ in *t* (Fig. 7B), the number of orderings is given by the second summand of the enumerator.

**Figure 7:**
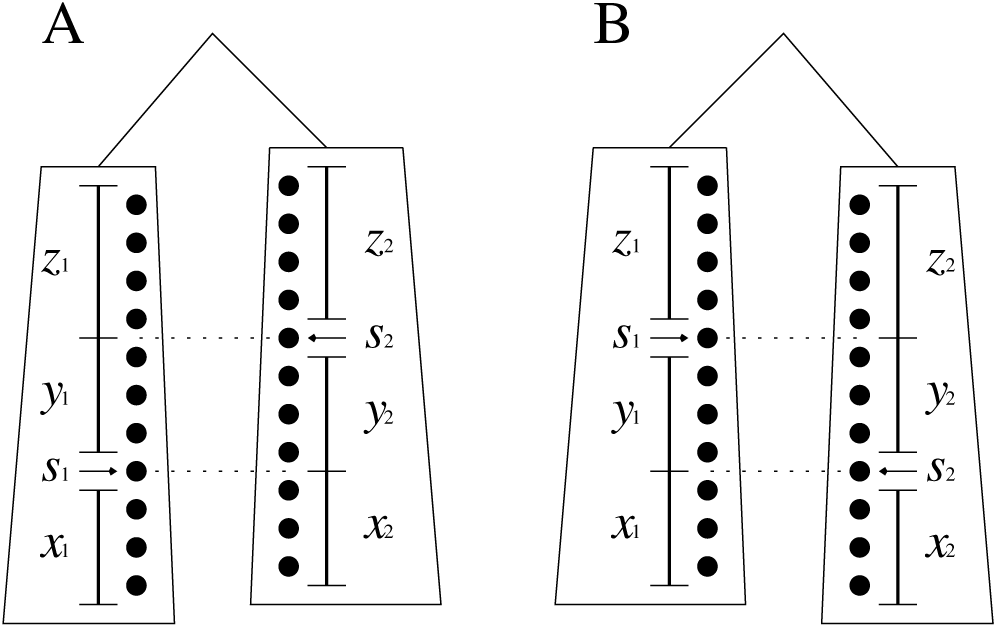
Tree decompositions for calculating Eq. (22).

For the case depicted in Fig. 7A,

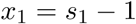

is the number of internal nodes of *t*_1_ with ranking smaller than *s*_1_. Similarly, *x*_2_ + *y*_2_ = *s*_2_ − 1 is the number of internal nodes of *t*_2_ with ranking smaller than *s*_2_. Also, we have *y*_1_ + *z*_1_ = *n*_1_ − 1 − *s*_1_ and

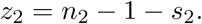

If *x*_2_ counts the nodes of *t*_2_ that in *t* are placed below the node *s*_1_ of *t*_1_, then 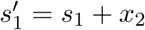. That is,

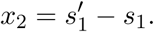

From the values of *x*_2_ + *y*_2_ and *x*_2_, we thus find

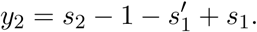

If *s*_1_ togehter with the nodes counted by *x*_1_ + *y*_1_ are placed below the node *s*_2_ in *t*, then 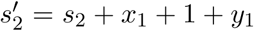. From the value of *x*_1_, we find

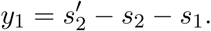

Finally, from the value of *y*_1_ + *z*_1_, we obtain

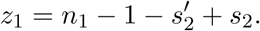

The number of orderings compatible with the values of 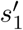 and 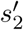 when we consider the decomposition of Fig. 7A is thus given by

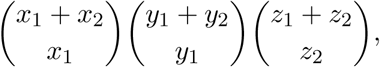

which is the first summand in the numerator of Eq. (22).

The decomposition of Fig. 7B, in which we consider the case of *s*_1_ placed above *s*_2_ in *t*, yields by similar calculations the second summand of the enumerator in Eq. (22). Observe that the two summands cannot be both different from zero at the same time. Indeed, the second binomial factor in the first summand is different from zero for 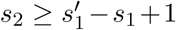, while the second binomial factor of the second summand is not zero for 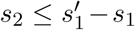.

By using the probability in Eq. (22), we can calculate *p*_2_(*s*|*s*_1_, *s*_2_, *n*_1_, *n*_2_) = Prob(*s*|*s*_1_, *s*_2_, *n*_1_ ≥ 2, *n*_2_ ≥ 2), that is, the conditional probability of *s* = *s* segments in the longest external branch of a random ordered history of size *n* = *n*_1_ + *n*_2_, given the number of segments *s*_1_ and *s*_2_ in the longest external branch of its rescaled root subtrees. Indeed, if *n*_1_, *n*_2_ ≥ 2, then the longest external branch of a history of size *n* = *n*_1_ + *n*_2_ is the longest external branch of its non-rescaled root subtrees. Thus, *p*_2_(*s*|*s*_1_, *s*_2_, *n*_1_, *n*_2_) can be written as

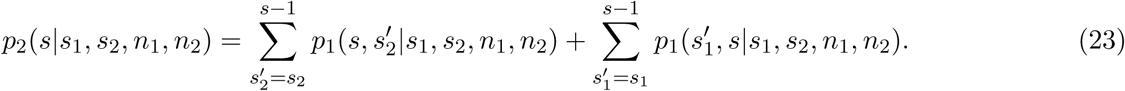

Finally, if *ω* ∈ [1, ⌊ *n/*2⌋] is the size of the smallest root subtree in a random Yule-distributed history (or uniformly distributed ordered history) of size *n*, then we can calculate the conditional probability *p*_*n*_(*s*|*ω*) ≡ Prob(*s*|*ω, n*) of *s′*= *s* segments in the longest external branch as

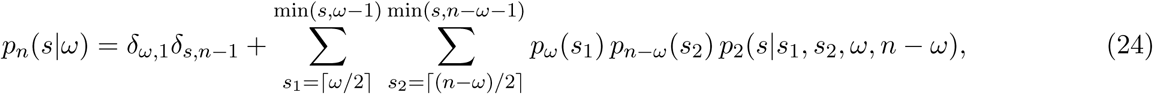

where *δ*_*s,n*−1_ is the probability of *s* segments when *ω* = 1, and *p*_*ω*_(*s*_1_) and *p*_*n*−*ω*_(*s*_2_) are respectively the probability of Eq. (17) that an ordered history of size *ω* and *n* − *ω* has *s*_1_ and *s*_2_ segments in the longest external branch. The probability in Eq. (24) is plotted for *ω* = 2, 3, 25 in Fig. 8 (left) for random Yule distributed histories of size *n* = 50. Interestingly, we observe that for *ω* = 3 and *ω* = 25 the probability of *s* segments in the longest external branch is basically the same. When *ω* = 2, the distribution is shifted to the left.

**Figure 8:**
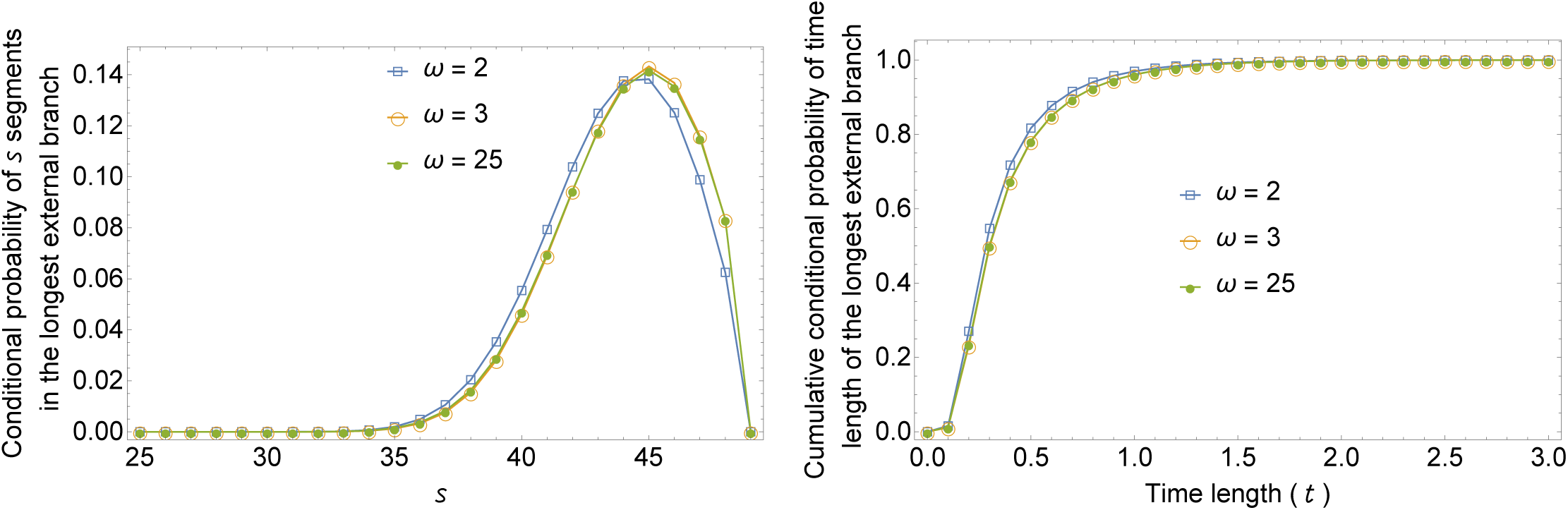
Left: Conditional probability that a Yule-distributed history of size *n* = 50 and *ω* = 2, 3, 25 has exactly *s* ∈ [⌈*n/*2⌉, *n* − 1] segments in its longest external branch (boxes for *ω* = 2, circles for *ω* = 3, and dots for *ω* = 25).Right: Cumulative conditional probability that the longest external branch of a history of size *n* = 50 and *ω* = 2, 3, 25 has length at most *t* (in coalescent time units).

By using Eqs. (2) and (24), the conditional probability that the time length *τ* of the longest external branch in a random history of size *n* with smaller root subtree of size *ω* lies in the interval *a < τ* ≤ *b* can be calculated as

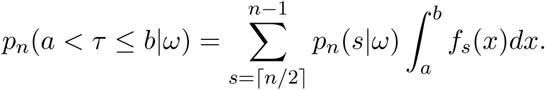

For *ω* = 2, 3, 25, the cumulative conditional probability *p*_*n*_(*τ* ≤ *t*|*ω*) is plotted in Fig. 8 (right) setting *n* = 50 and letting *t* range over the interval [0, 3] in steps of 0.1. As already observed for the number of segments, for *ω* = 3 and *ω* = 25 the values of *p*_*n*_(*τ* ≤ *t*|*ω*) are basically the same. When *t* ≤ 1, the cumulative conditional probability is larger for *ω* = 2 than for *ω* = 3, 25.

### 5.2 The number of segments in the second longest external branch

In Section 5.1, we have studied the random variable *s′* counting the number of segments in the longest external branch of a random history of given size. Note that for a history *t* there could be two external branches of size *s′* (*t*), when these branches form a cherry in *t*. Considering the set of external branches of *t* that are strictly shorter than the longest one(s), we define *s″* (*t*) *∈* [⌊ (*n* − 1)*/*2⌋, *s′* (*t*) − 1] to be the number of segments in the longest external branch of this set, and we say that *s* (*t*) is the number of segments of the second longest external branch of *t*. For example, *s″*(*t*) = 4 when *t* is the history depicted in Figure 1. In this section, we study the distribution of the random variable *s″* (*t*), when *t* is a random history of size *n* selected under the Yule probability model. By using the equivalence with the uniform distribution over ordered histories of size *n* or permutations of size *n* − 1, we find that the probability of *s″* = *s* segments in the second longest external branch of a random history of size *n* can be expressed as a simple function of the probability that a random history of size smaller than *n* has *s′* = *s* segments in its longest external branch.

We first calculate the joint probability of *s′* and *s″*. Let Π_*n*_(*X*) denote as in section 5.1 the number of permutations of size *n* with peak entries matching the elements of the set *X* ⊆ [3, *n*], and choose *s*_1_ and *s*_2_ such that ⌉*n/*2⌋ ≤ *s*_2_ *< s*_1_ ≤ *n*. For a fixed set *Z* ⊆ [3, *s*_2_ − 1], by setting *S* = *Z* ∪ [*s*_2_ + 1, *s*_1_ − 1] and *k* = *n* − *s*_1_ in Lemma 3.3 of [4], for *n* ≥ 2 we find

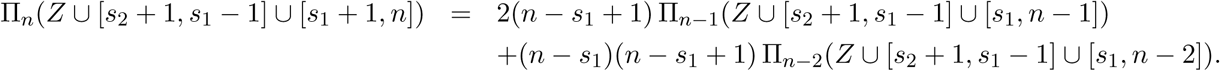

If we sum both sides of the latter equation over the possible subsets *Z* of [3, *s*_2_ − 1], then we obtain

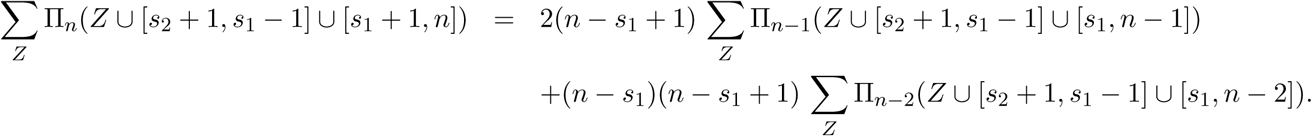

where the sum ∑_*z*_ ∏_*n*_(*Z*∪[*s*_*2*_+1, *s*_1_ − 1] ∪[*s*_*1*_+1,*n*]) counts the permutations of size *n* in which the largest and the second largest non-peak entry are respectively *s*_1_ and *s*_2_, and the sums ∑_*z*_ ∏_*n*−1_(*Z*∪ [*s*_*2*_+1, *s*_1_−1] ∪[*s*_*1*_, *n*−1]) and ∑_*z*_ ∏_*n*−2_(*Z*∪ [*s*_*2*_+1, *s*_1_−1] ∪[*s*_*1*_, *n*−2]) count respectively the permutations of size *n* − 1 and *n* − 2 in which the largest non-peak entry is *s*_2._ For instance, let us set n = 6. For *s*_1_ = 5 and *s*_2_ = 4, Table 2 of [4] gives ∑_*z*_ ∏_*n*_(*Z* ∪ [*s*_*2*_ + 1, *s*_1_ − 1] ∪ [*s*_*1*_ + 1,*n*]) =264, ∑_*z*_ ∏_*n*−1_(*Z* ∪ [*s*_*2*_ + 1, *s*_1_ − 1] ∪ [*s*_*1*_, *n* − 1]) = 60 and ∑_*z*_ ∏_*n*−2_(*Z*∪ [*s*_*2*_+1, *s*_1_−1] ∪[*s*_*1*_, *n*−2]) = 12, where 264 = 2 · 2 · 60 + 1 · 2 · 12.

By using the correspondence between non-peak entries of permutations of size *n* and external branches of ordered histories of size *n* + 1 (Section 2), the number 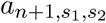 of ordered histories *t* of size *n* + 1 in which *s*′(*t*) = *s*_1_ and *s*″(*t*) = *s*_2_ can be calculated as

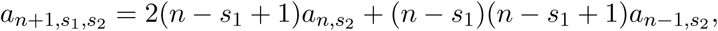

where 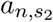 and 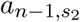 count the number of ordered histories of size *n* and *n* − 1, respectively, in which the longest external branch has *s*_2_ segments. Replacing *n* + 1 by *n* in the latter equation and dividing by the number (*n* − 1)! of ordered histories of size *n*, the probabilites *p*_*n*−1_(*s*_2_) and *p*_*n*−2_(*s*_2_) of Eq. (17) yield the joint probability

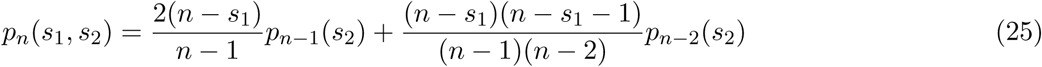

of a random history of size *n* in which the longest external branch has *s*′ = *s*_1_ segments and the second longest external branch has *s*″ = *s*_2_ segments. The conditional probability of *s*″ = *s*_2_ segments in the second longest external branch given *s*′ = *s*_1_ segments in the longest external branch of a random history of size *n* is thus 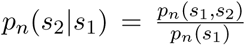 (Fig. 9, left). The sum 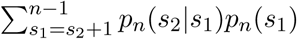 gives the unconditional probability of *s*″ = *s*_2_ segments in the second longest external branch of a random history with *n* leaves (Fig. 9, right).

**Figure 9:**
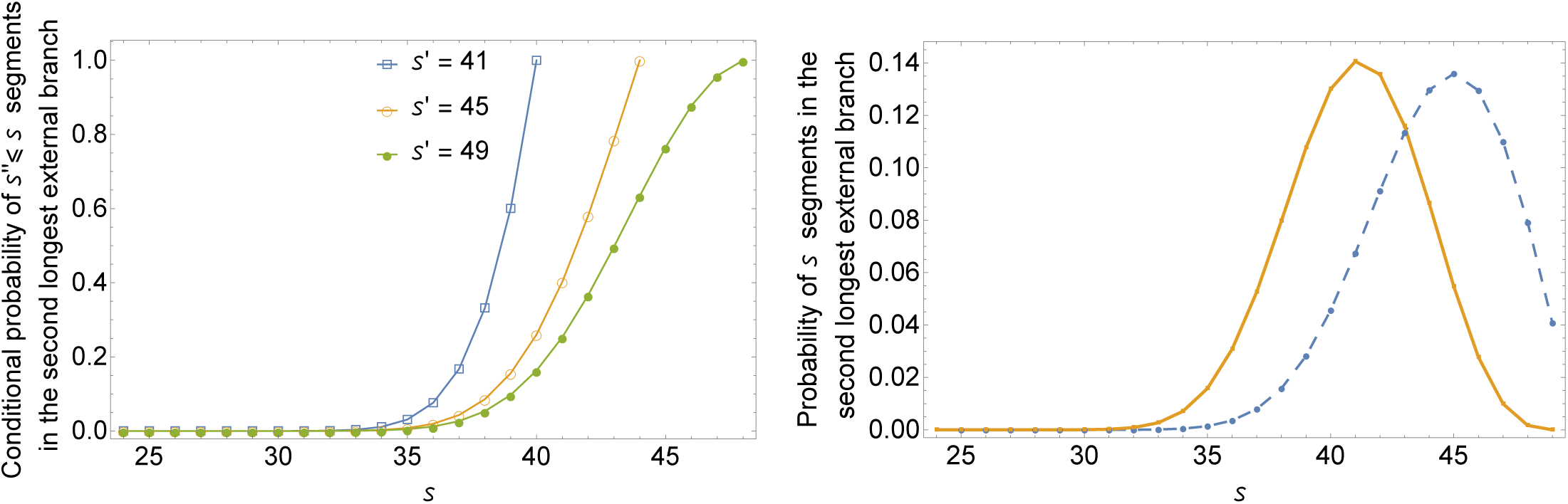
Left: Conditional probability of *s″* ≤ *s* segments in the second longest external branch of a random Yule-distributed history of size *n* = 50, when the longest external branch has *s′* = 41, 45, 49 segments (left to right curve). Right: Unconditional probability that a random Yule-distributed history of size *n* = 50 has exactly *s* segments in its first (dashed line) and second (solid line) longest external branch.

**Figure 10:**
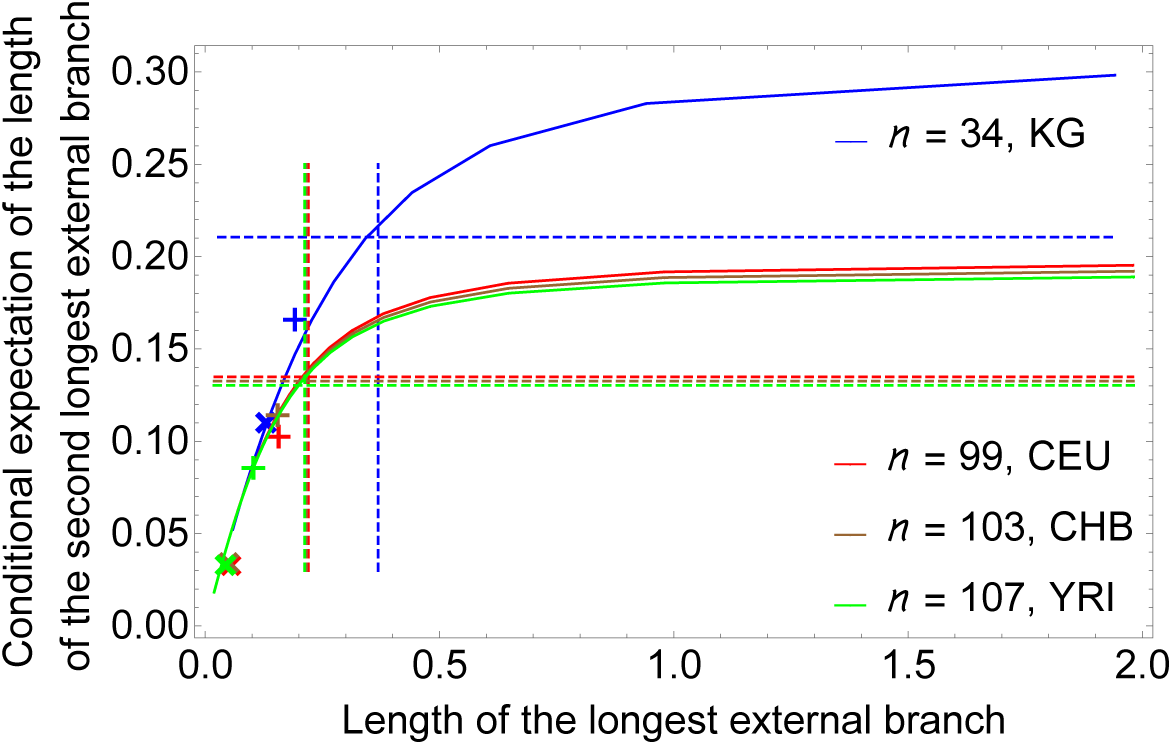
Estimates of the longest and second-longest external branches in coalescent trees. Red curves and symbols: mitochondrial data from *Homo sapiens*, CEU population (Central European origin); green: YRI population (Yoruba in Nigeria); brown: CHB population (Han-Chinese from Beijing). Blue curves and symbols: nuclear data from *Danio rerio*, CB population (Cooch Behar, India). *n*: sample size (# of chromosomes examined); +-symbol: estimate *E*_1_ (using all available polymorphisms); ×-symbol: estimate *E*_2_ (using only singletons). Solid lines: conditional expectation of the length of the second-longest branch, given the length of the longest branch. For a fixed value of *n*, every integer *s* ∈ [⌊ *n/*2 ⌋, *n* − 1] yields a point (*x*_*s*_, *y*_*s*_) of the corresponding theoretical line. The abscissa *x*_*s*_ is the expected time length of an external branch with *s* segments (1). The ordinate *y*_*s*_ is the expected time length of the second longest external branch, when the first longest has *s* segments (Section 5.2). Dashed lines: average length of the longest (vertical) and (un-conditioned) average length of the second-longest branch (horizontal). Refer to Section 6 for more detailed description.

## 6 Estimating the longest external branches from experimental data

To compare the theoretical results obtained above with experimental data we estimate the longest external branches of a hypothetical coalescent tree from the maximal counts of singleton mutations in sets of SNP data. Since recombination can disturb tree topology, we concentrate on (i) the non-recombining mitochondrial genomes from *Homo sapiens* (size about 17kb) from three different populations (CEU, CHB and YRI) and (ii) on a short 7.3kb genomic sequence (pre-mRNA) of a single nuclear gene (CTCF) from a wild population of zebrafish. Human data (1*k* genomes initiative, phase III) were downloaded from www.1000genomes.org. The zebrafish sample comes from a larger collection of data, which we sequenced and analyzed as part of a different project [20]. The sample analyzed here has a size of *n* = 34, from 17 individuals collected from the wild (GPS coordinates N022.262 E087.279; sub-population termed ‘KG’).

We calculated two simple estimates for the relative length of the longest and second longest branches of a coalescent tree: *E*_1_ is based on the total number of SNPs observed (*S*_total_), *E*_2_ is based on the observed singletons only (*S*_singl_). Since the combined length of all external branches compares to the total tree length as 1 to *h*_*n*−1_ [21], we estimate

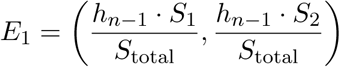

and

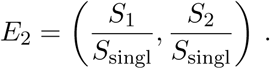

The observed and estimated data are collected in Table 1. All data fit very well the theoretical prediction. For all populations both coordinates of estimate *E*_1_ are larger than those of *E*_2_. This is compatible with the notion that purifying selection leads to an increase of singleton mutations compared to the neutral expectation. However, purifying selection affects all individuals in the same way, hence is not inducing a bias on the longest or second longest branch. The larger values of Danio compared to human are explained by different sample sizes (also visible in the shift among the solid curves representing the theoretical values). Comparing the derived human populations (CEU and CHB) to the ancestral African population (YRI), one observes a slight increase in both coordinates of *E*_1_ in CEU and CHB compared to YRI. This is explained by the well known stronger increase in the number of singletons in the frequency spectrum of the derived populations due to the bottleneck effect accompanying the migration out of Africa.

**Table 1:**
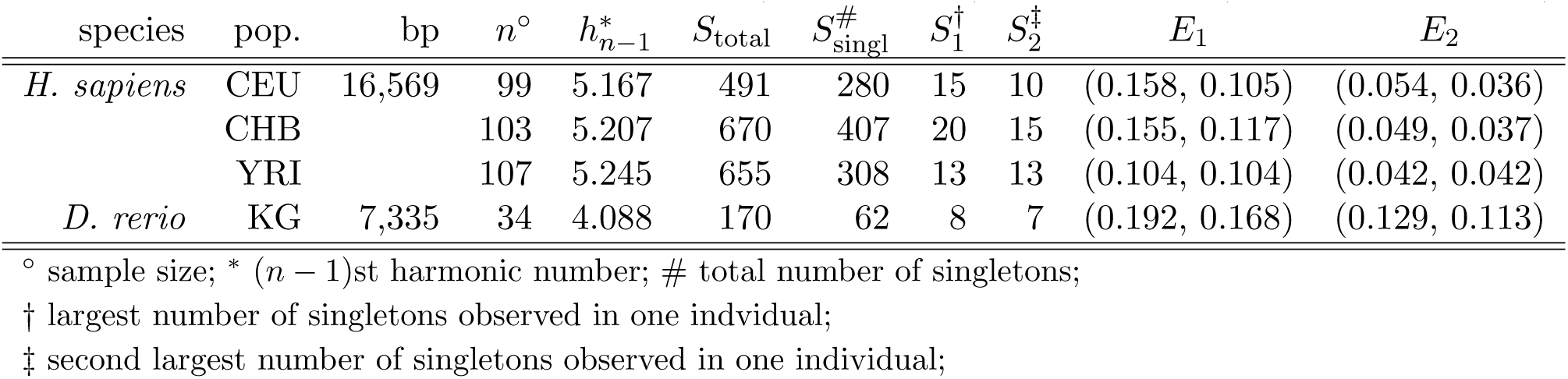
Experimental data observed in human (mitochondria) and zebrafish (genome-wide survey).

| species | pop. | bp | $n^\circ$ | $h_{n-1}^*$ | $S_{\text{total}}$ | $S_{\text{singl}}^\#$ | $S_1^\dagger$ | $S_2^\ddagger$ | $E_1$ | $E_2$ |
| --- | --- | --- | --- | --- | --- | --- | --- | --- | --- | --- |
| *H. sapiens* | CEU | 16,569 | 99 | 5.167 | 491 | 280 | 15 | 10 | (0.158, 0.105) | (0.054, 0.036) |
|  | CHB |  | 103 | 5.207 | 670 | 407 | 20 | 15 | (0.155, 0.117) | (0.049, 0.037) |
|  | YRI |  | 107 | 5.245 | 655 | 308 | 13 | 13 | (0.104, 0.104) | (0.042, 0.042) |
| *D. rerio* | KG | 7,335 | 34 | 4.088 | 170 | 62 | 8 | 7 | (0.192, 0.168) | (0.129, 0.113) |

$^\circ$ sample size; $^*$ $(n - 1)$ st harmonic number; $^\#$ total number of singletons;
$^\dagger$ largest number of singletons observed in one individual;
$^\ddagger$ second largest number of singletons observed in one individual;

## 7 Conclusions

We have studied probabilistic properties of the number of segments present in the external branches of a random Yule-distributed history of given size. The approach followed in our calculations also provides a combinatorial framework for the analysis of the time length of the external branches of a Kingman coalescent tree of given size.

In Section 3, we have focused on the probability that a random history of fixed size has a given number of external branches of a given number of segments. Eq. (3) together with Eqs. (4-10) enable respectively the calculation of the unconditional and conditional probability of *µ* ∈ {0, 1, 2} external branches of *s* ∈ [1, *n* − 1] segments in a random history of size *n*. The unconditional probability of *µ* = 0 (resp. *µ* = 2) external branches of *s* segments increases (resp. decreases) for increasing values of *s*. In particular, it is interesting to observe that the probability of missing an external branch of *s* segments symmetrically corresponds to the probability of having two external branches of *n* − *s* + 1 segments. Also, in a random history, the probability of exactly one branch of *s* segments is equal to the probability of exactly one branch of *n* − *s* + 1 segments (Fig. 2). Numerical plots of the conditional probability of *µ* external branches of *s* segments given *µ*_*r*_ external branches of *r* segments are given in Fig. 3.

In Section 4, we have used known combinatorial results [4] on the set of peak entries of a permutation of size *n* − 1, for characterizing the possible sets of missing external branches in a history of size *n*. Furthermore, for a given subset 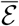, Eq. (12) provides a recursive formula for calculating the probability that the external branches missing in a random history of size *n* are exactly those whose number of segments is listed in the set 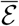 (Fig. 4).

In Section 5, we have studied the probability of a given number of segments in the longest and second longest external branch of a history of size *n* selected at random. The first probability can be evaluated through Eqs. (16, 17). The joint probability is given by Eq. (25). In Section 5.1.2, we have derived Eqs. (22-24) for computing the conditional probability of the number of segments in the longest external branch of a random history of *n* leaves given the size *ω* ∈ [1, ⌊*n/*2⌋] of its smallest root subtree. Investigations of Section 5 yielded two findings on the structure of Yule histories that were not intuitively clear before. Surprisingly, imbalance does (almost) not influence the length of the longest external branch. For sufficiently large *n*, the distribution of the length of the longest external branch is roughly the same when we condition on different values of *ω ≥* 3 (Fig. 8). Furthermore, we observe that conditioning on the discrete length *s*′ of the longest external branch can strongly affect the distribution of the number of segments *s*″ in the second longest external branch. In Fig. 9 (left), we see that a relative small decrease of the value of *s*′ from *s*′ = 49 to *s*′ = 41 results in a step function behavior of the conditional probability of *s*″ being smaller than a given value *s*.

The study of the number of segments in the external branches of a random Yule-distributed history can assist in the analysis of the time length of the external branches of a Kingman coalescent tree, which can be seen as Yule-distributed history in which the time length of the *i*th layer is an exponentially distributed variable. Importantly, the probability density function *f*_*s*_(*x*) of the time length of an external branch of *s* segments in a history of size *n* can be evaluated as in Eq. (2), and our study of the discrete length of external branches be used as a benchmark for biological data scenarios. In particular, when analyzing the mutation frequency spectrum of population samples, one may be interested in the question whether the number of singletons seen in a single haplotype significantly exceeds neutral expectation. For instance, this could be an indication of unaccounted population substructure. In Section 6, we applied our theoretical results on the length of the longest and second longest external branch to sequence samples from human and from a wild population of zebrafish. For the latter, an initial suspicion of sample contamination with non-genuine material was not confirmed with our results.

Several direction of research naturally arise from our work. First, it would be of interest to study the random variables considered in this article under different probability models—e.g., assuming a uniform distribution over the set of unordered histories of given size. Histories with a larger number of cherries have a smaller probability to be generated under the Yule process. Switching to a different distribution could for instance affect the probability of external branches of multiplicity 2 or the correlation between root imbalance and length of the longest external branch. Second, it would be important to extend the approach used in this article for studying the length of the external branches of a Yule-distributed history or coalescent tree to consider also the length of the branches ancestral to a cherry, which are associated with doubletons in the mutation frequency spectrum. Finally, we observe that encoding the ordered histories of size *n* as permutations of size *n* − 1 allows to define a geometric structure over the set of histories of size *n*, when these are grouped together in equivalence classes according to their set of external branches. In particular, the admissible sets of missing external branches of the histories of size *n*, that is, the possible peak sets of the permutations of size *n* − 1, are shown in [4] to form an abstract simplicial complex over the vertex set [3, *n* − 1]. It seems natural to ask for possible biological interpretations of this complex.

## Acknowledgments

Support was provided by a Rita Levi-Montalcini grant to FD from the Ministero dell’Istruzione, dell’Università e della Ricerca and by a grant from the German Research Foundation (DFG SPP-1590) to TW.

## Appendix Proof of Eq. (15)

The formula in Eq. (15) can be rewritten as 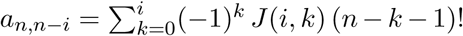, where 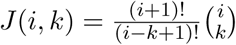. Setting *ã*_*n,i*_ = *a*_*n,n*−*i*_, Eq. (15) is thus equivalent to

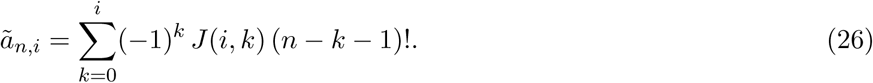

From 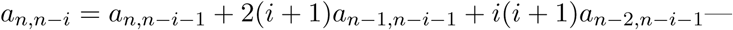 which is Eq. (14) with *s* = *n* − *i*—we obtain the recurrence 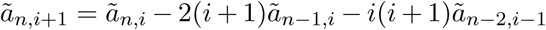. Replacing *i* + 1 by *i* in the latter expression yields

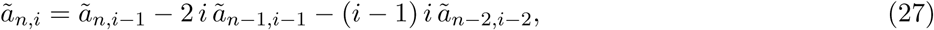

which we use to show formula (26) by induction on *i*. Substituting (26) in the right-hand side of (27), we find

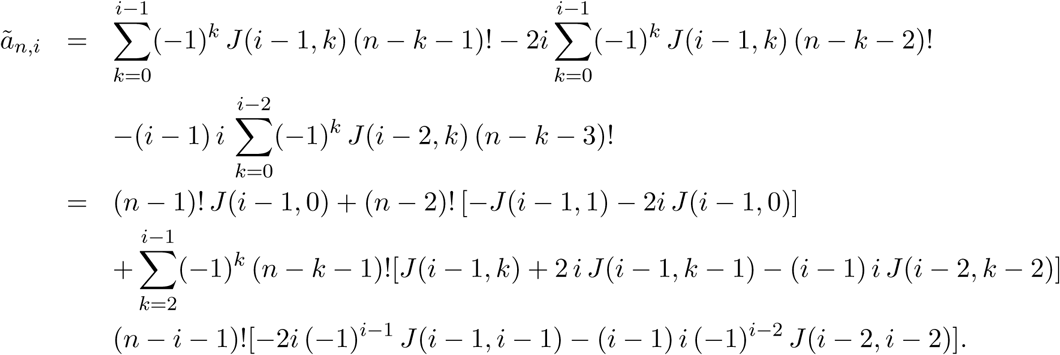

For 0 *≤ k ≤ i*, the coefficients of (*n* − *k* − 1)! in the latter expression can be easily seen to satisfy

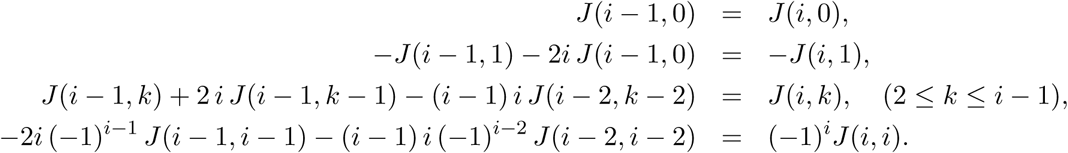

Hence, the formula obtained recursively for *ã*_*n,i*_ matches the sum in Eq. (26).

